# “Antibiotic inhibition of bacteria growth in droplets reveals heteroresistance pattern at the single cell level”

**DOI:** 10.1101/328393

**Authors:** Ott Scheler, Karol Makuch, Pawel R. Debski, Michal Horka, Artur Ruszczak, Natalia Pacocha, Krzysztof Sozański, Olli-Pekka Smolander, Witold Postek, Piotr Garstecki

## Abstract

Heteroresistance is a phenomenon where isogenic bacteria population exhibits a diverse antibiotic resistance pattern at sub-population or single cell level. The sub-populations with higher resistance can remain undetected with conventional diagnostics which makes them subsequently harder to treat. Such surviving phenotypically heterogeneous sub-populations are also a potential hotbed for novel mutations, thus increasing the resistance permanently in bacteria. Droplet microfluidics gives tools for high-throughput analysis of bacteria and their response to antibiotics at single cell level, which is difficult to obtain with traditional agar plate technologies. In here we show for the first time the precise digital quantification of drug resistance profile in isogenic population at single cell level. We also see that the inhibiting amount of drug per bacteria remains quite stable regardless of bacteria density. Interestingly, the bacteria clump together preferably near these sub-inhibitory conditions. The technology and findings we describe here provide novel quantitative insight into the heteroresistance which is a key step in understanding the pathways leading to drug resistance. This knowledge is crucial in the context of global drug resistance threat as it can help us to find tools to prevent further escalation of drug resistance.

## Introduction

- Heteroresistance is a phenomenon where isogenic bacteria population exhibits a diverse antibiotic resistance pattern at sub-population or single cell level^1^.
- The subpopulations with higher resistance can remain undetected with conventional diagnostics which makes them subsequently harder to treat^2^.
- Such surviving phenotypically heterogeneous sub-populations are also a potential hotbed for novel mutations, thus increasing the resistance permanently in bacteria^3^.
- Droplet microfluidics^4^ gives tools for high-throughput analysis of bacteria and their response to antibiotics at single cell level^5^, which is difficult to obtain with traditional agar plate technologies^1^.
- In here we show for the first time the precise digital quantification of drug resistance profile in isogenic population at single cell level.
- We also see that the inhibiting amount of drug per bacteria remains quite stable regardless of bacteria density.
- Interestingly, the bacteria clump together preferably near these sub-inhibitory conditions.
- The technology and findings we describe here provide novel quantitative insight into the heteroresistance which is a key step in understanding the pathways leading to drug resistance. This knowledge is crucial in the context of global drug resistance threat^6,7^ as it can help us to find tools to prevent further escalation of drug resistance.

## Main text

> [In gram-negative bacteria the constant evolution of beta-lactamases is increasing the spread of resistance against beta-lactam antibiotics]

Beta-lactamases function by degrading beta-lactam antibiotics that inhibit bacteria growth by disrupting cell wall synthesis^8,9^. One of the most widespread beta-lactamase protein families associated with resistance is TEM-family^8,10^. Here we investigate the cefotaxime resistance profile of TEM-20 beta-lactamase in model strain *Escherichia coli* Dh5α at the single cell level. The strain carries plasmid with TEM-20 gene and a second plasmid with yellow fluorescent protein (YFP) for detection. We use minimum inhibitory concentration (MIC) of a drug that prevents bacteria proliferation as a measurement of resistance^11^.

> [For capturing isolated single-cell growth response pattern we encapsulate bacteria into water-in-oil droplets]

We used microfluidic chip with flow-focusing geometry to generate libraries of monodisperse 2nL droplets that each act as a separate miniature “test-tube” (Fig1A). We encapsulate cells with antibiotic (separate droplet library for each cefotaxime concentration), incubate bacteria overnight and screen the droplets for increased fluorescence from fully grown (saturated) colonies. The term “colony” here stands for an accumulation of microbes in droplet, usually occurring as a clone of a single original organism^4,12^. Therefore, the single cell in this article is equivalent to single colony forming unit (CFU). One bacteria per 2nL droplet is equivalent to 5×10^5^ CFU/mL which is the standard for MIC tests^11,13^. In principle, the antibiotic screening in droplets is ‘digital MIC assay’^14,15^ where outcome of each discrete experiment in droplet is binary: ‘1-positive’ with detectable bacteria growth or ‘0-negative’ without growth. For single cell analysis it is necessary to avoid encapsulating two or more bacteria as much as possible. In order to achieve that most of the generated droplets should be empty and not contain any bacteria because encapsulation events in such systems are described by the Poisson probability distribution. Thus, for single cell analysis the encapsulation rate *λ* is recommended to be 0.1-0.3 or lower^15^. In our case the *λ* was ∼0.18 as measured from control experiment without antibiotic. For each antibiotic concentration we screened libraries containing ∼1500 droplets with bacteria.

**Figure 1:**
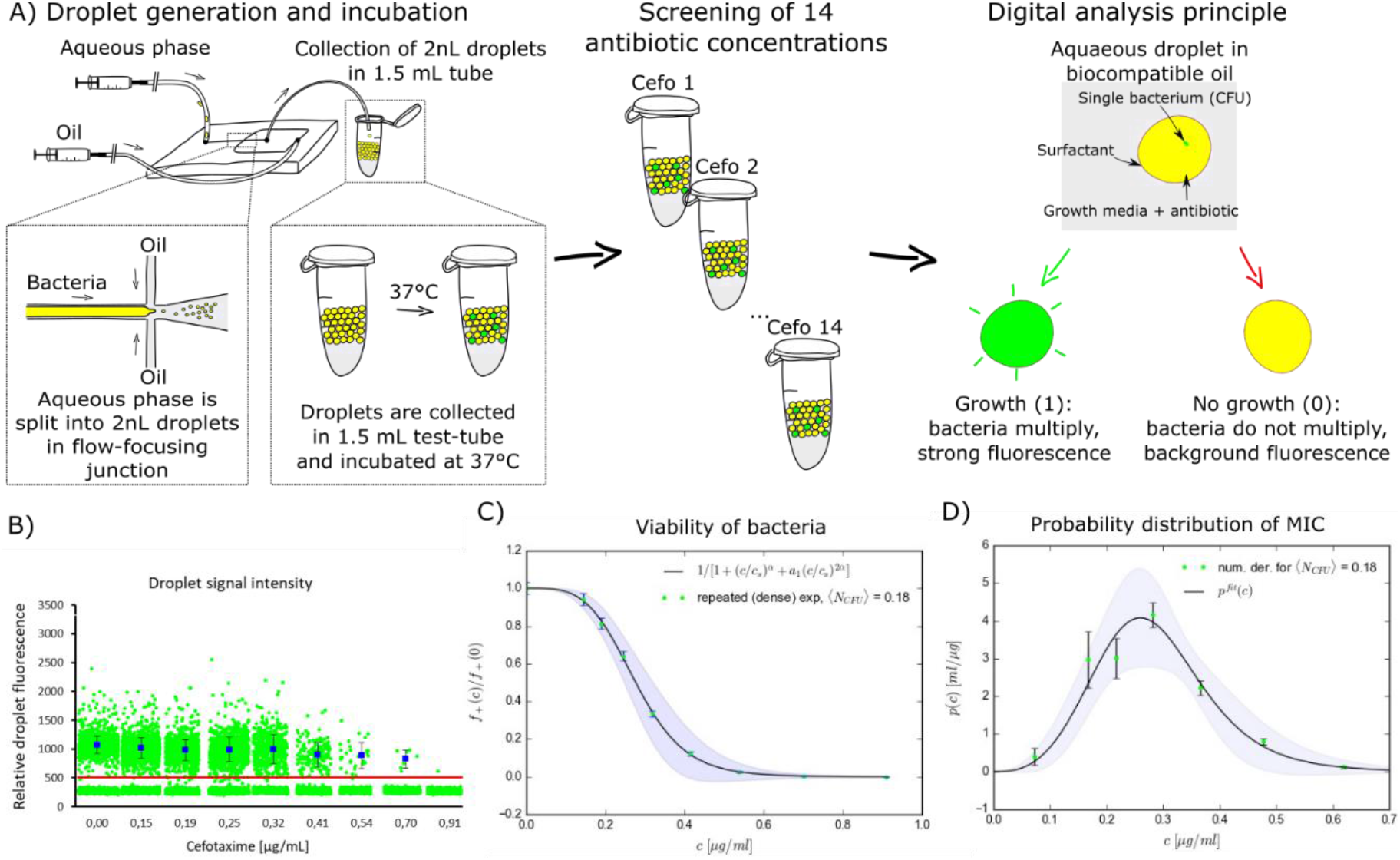
Cefotaxime reveals heteroresistance pattern at single cell level. A) Schematic of the single cell droplet assay where aqueous phase (consisting of bacteria, media and antibiotics) are encapsulated into water-in-oil-droplets using HFE 7500 with 2 % surfactant as a continuous phase. Each different antibiotic concentration was screened in separate library. The outcome of the bacteria assay in droplet is ‘digital’ in principle: the bacteria either grows (1) or not (0). B) Signal intensities of each droplet in the experiment. Red line at 500 marks the threshold for positive droplets. Blue rectangles show the average signal of positive droplets with standard deviation as error bars. C) Fraction of positive droplets normalized by its value for experiment without antibiotic, *f*_+_ (*c*)/*f*_+_ (0) as a function of antibiotic concentration *c*. Continuous line represents the following fit 1/[1 + (*c*/*c_s_*)^*α*^ + *a*_1_(*c*/*c_s_*)^2*α*^] with the fitting parameters *c_s_* = 0.29*μg*/*ml, α* = 3.78, *a*_1_ = 0.26 determined by the least square method. D) Points: probability distribution of individual MIC in the population obtained by numerical derivative from the data points in Fig. 1C (see Supplementary part 7 for further explanation). Continuous line represents the derivative of the fit from Fig. 1C

> [We saw that the isogenic bacteria population has a diverse resistance profile at single cell level]

The response curve of growth inhibition by cefotaxime decreases gradually with antibiotic concentration increase and there is no sharp (Heaviside step function) transition between “growth” and “no growth” response (Fig 1C). At the same time the colony density (fluorescence intensity) in positive droplets remains on the same level even when the fraction of positive droplets significantly drops (Fig 1B). This observation suggests, that proliferation in droplets is a binary stochastic variable – the individual cells either grow into colonies or not.

The heterogeneity of the whole population is characterized by the probability distribution density, *p*(*c*), where growth of a single bacterium in a droplet is inhibited by the antibiotic concentration *c*. The probability distribution is defined for the single bacteria in a droplet. This assumption is achieved when there is a big number of empty droplets, *f*_+_(0) ≪ 1. Then the fraction of resistant bacteria in the population, 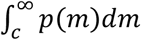, is given by, 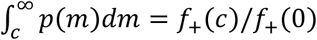. The normalized fraction of positive droplets, *f*_+_(*c*)/*f*_+_(0), is shown if Fig. 1C with errors obtained as described in Supplementary part 5. In principle, such probability was suggested recently by Lyu *et al*.^5^, but the pattern and full diversity of the heteroresistance is described here for the first time.

Such resistance profile qualifies as a heteroresistance by definition as the measured MIC increases within the population by more than 8-fold^1^. Emergence of spontaneous resistant mutants is highly unlikely to explain the breadth of this distribution as the mutation rate in bacteria is orders of magnitude lower^16^ than the fraction of growing bacteria in this experiment. This observation cannot be explained by the presence of persister cells either as the bacteria grow under constant antibiotic exposure in our setup^17^.

> [The heteroresistance is most likely caused by differences specifically in TEM-20 expression level or more globally on the whole transcriptome profile in the cells]

In our model system the TEM-20 gene expression is modulated by at least two components: i) the copy-number of TEM-20 carrying plasmid in the cell and ii) general stochasticity of the gene expression^18,19^.

> [We looked how susceptibility to the drug changes with increasing bacteria inoculum density]

β-lactam antibiotics are often subject to the inoculum effect (IE) where the efficiency of the drug depends on the starting inoculum density of the bacteria^20^. IE is often overlooked in traditional MIC-assays due to the inaccuracy in setting inoculum density to recommended level using conventional OD-measurement^13^.

> [For precise estimation of bacteria densities and measuring IE we used color-coded virtual array of droplets]

We use “virtual array” strategy described by Abate *et al.^21^* to pool together droplet libraries for easier downstream handling and analysis. We prepared 16 two-fold dilutions of the *E. coli* and labelled each dilution with combination of two fluorescent dyes: Cascade Blue and Alexa 647 (Fig 2A-C). Immediately after generation the 16 color-coded libraries were pooled together into single 1.5 μL test-tube for further overnight incubation at 37 °C.

**Figure 2:**
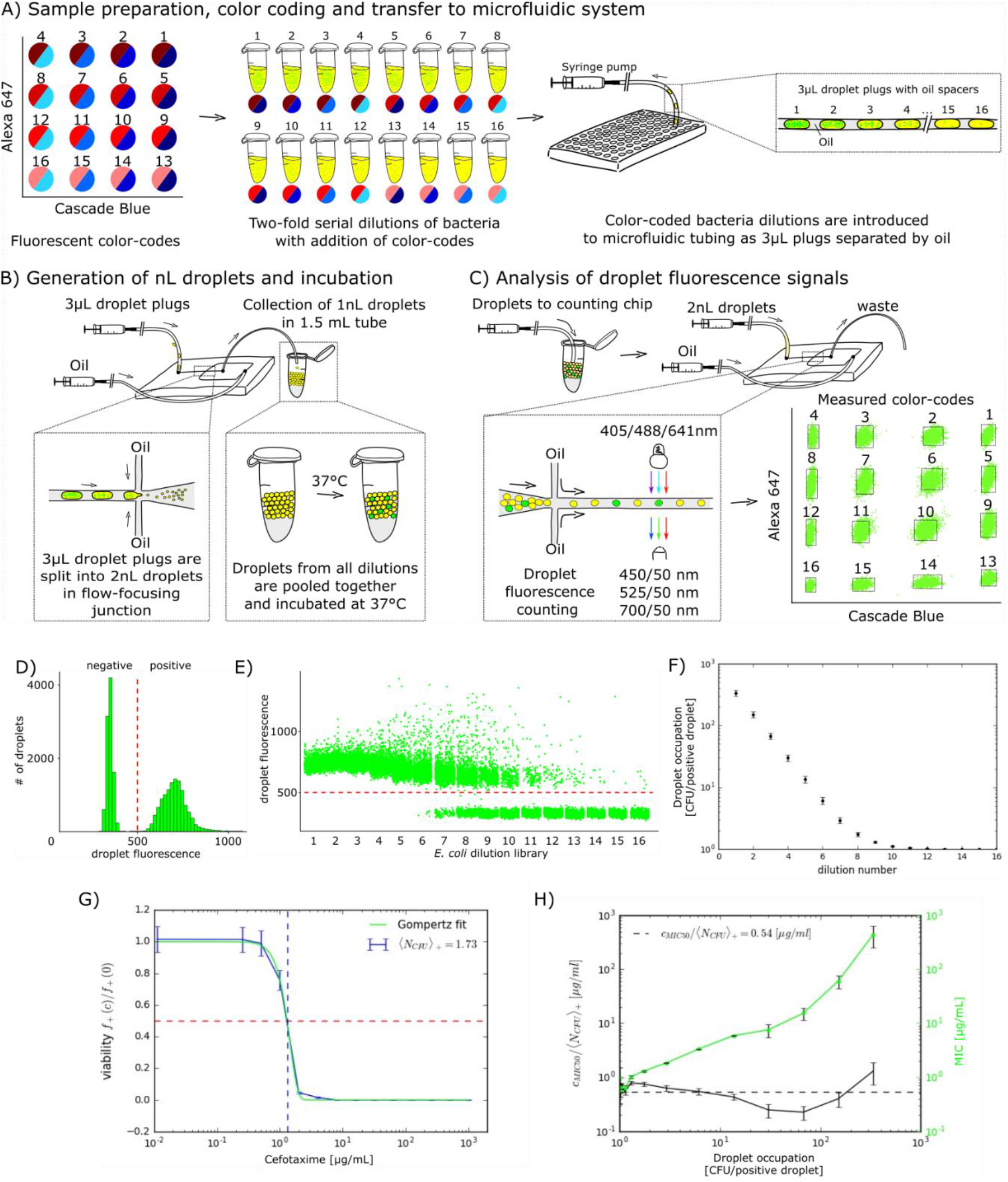
inhibiting amount of cefotaxime per bacteria remains stable over wide range of bacteria densities. A) a schematic for color-coding different bacteria densities with Cascade Blue and Alexa 647 fluorescent dyes. Both dyes are represented in the virtual array as 4×4 matrix of 16 color-code combinations. Two-fold serial dilutions of bacteria are color-coded and introduced to microfluidic tubing as oil-separated 3 μl plugs. B) Generation of “virtual array” from color-coded bacteria densities. Plugs are split into 2 nL droplets that are pooled together into single library. C) Droplet fluorescence measurement in three different channels. The virtual array is inserted into the counting chip that is fixed in a confocal microscope stage to measure the intensity of fluorescence emitted by droplets. For spacing the droplets during the measurement, an additional stream of carrier oil is introduced into the flow-focusing junction immediately before the acquisition area. Each droplet is then given “the address” in the virtual array based on its gating into the bounding box. D) Histogram of the pooled droplet signals showing bacteria growth in green channel. Red dashed line notes the threshold between negative and positive droplets. E) Plot showing the bacteria growth (green channel) separately in each droplet. On X-axis the droplets are sorted according to their color-code allocation in the virtual array (the same data as in Fig 2D). Note: There is significant population of droplets with high fluorescence intensity near value 1000 and above that are explained in “clumping” section. F) Graph showing the average droplet encapsulation rate of bacteria in different virtual array libraries (NCFU+). G) Calculation of MIC in the library with bacteria density 1.73 NCFU+. Green line show the Gompertz fit of the data and red dashed line the 0.5 viability fraction. Blue vertical dash shows the position of the MIC value. H) Graph showing comparison of MIC (green) and MIA (black) over different inoculum densities.

> [We calculated initial inoculum density in virtual array with digital counting algorithm and equations^14,22^]

We set a threshold to distinguish between positive and negative droplets based on their fluorescence (Fig 3D). Next we investigated separately each library in virtual array to calculate their positive droplet fractions (Fig 3E). We calculated the average encapsulation event for each library as “CFU/positive bacteria containing droplet” (Fig 3F). See Supplementary part 6 for more detailed explanation of calculations. The average CFU/droplet ranged over two orders of magnitude from 1 to ∼338 per droplet. That translates to inoculum range between 5×10^5^ and ∼1.7× 10^8^ CFU/mL by conventional approach.

**Figure 3:**
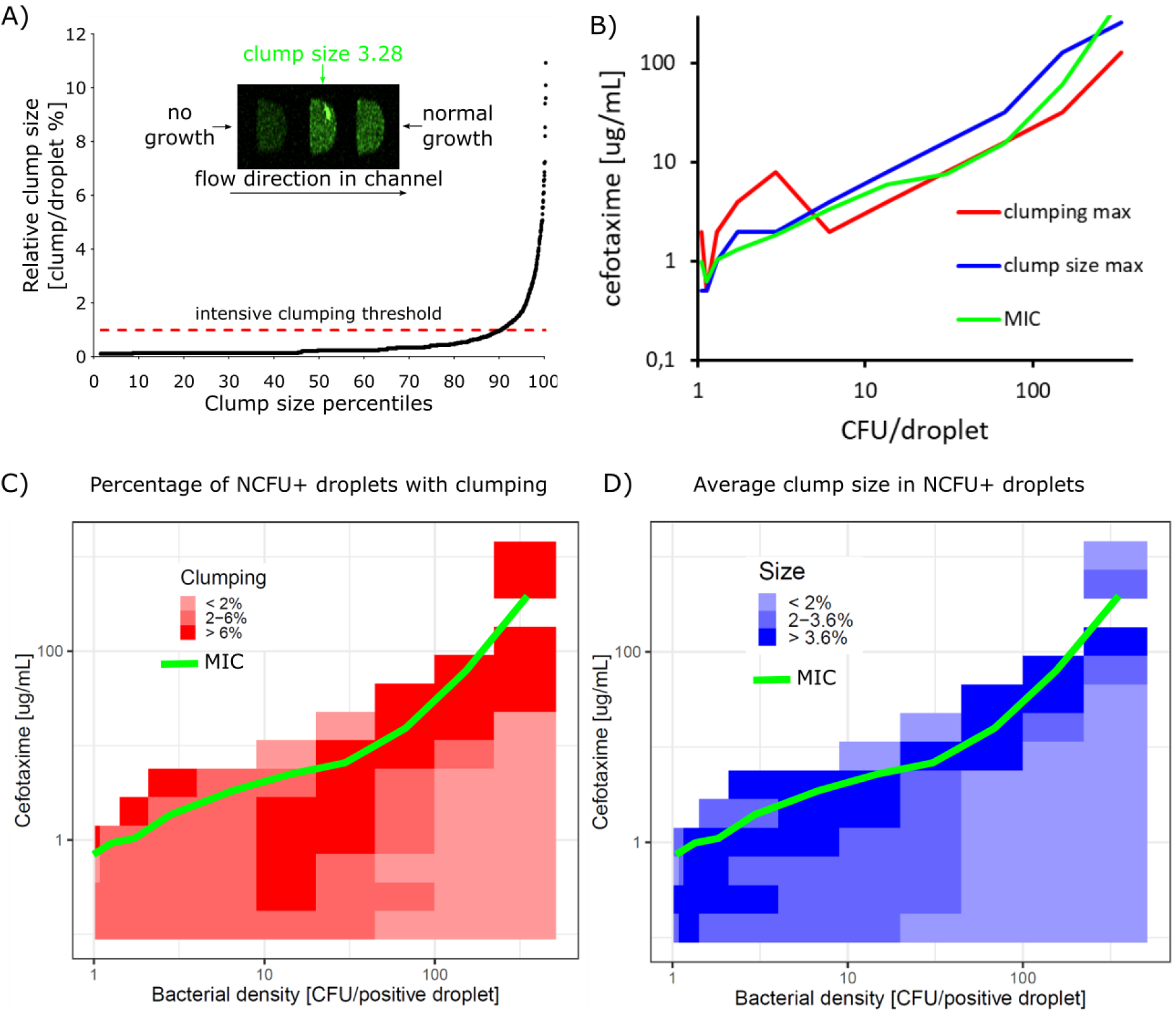
cefotaxime modulates the clumping of bacteria. A) Intensive clumping threshold is set based on the relative clump size in droplets. We consider the 90^th^ percentile of clump sizes to have intensive clumping covering at least ∼1% of droplet area. B) At different inoculum densities both the highest clumping rate (red line) and the greatest size of clumps (blue line) occur near sub-inhibitory cefotaxime conditions (green line). C) and D) heat-maps showing the relative clumping rates (red) and clump sizes (blue) in the matrix of different bacteria densities (X-axis) and cefotaxime concentrations (Y-axis). Green line shows the approximate position of MIC in these experiments

> [We investigated IE in two-fold dilution series of cefotaxime ranging from 0.25 to 1024 μg/mL]

Each antibiotic concentration is analyzed in separate virtual array. We use virtual array without antibiotics as a control to show bacteria growth without antibiotic stress.

> [In this paper we calculate the MIC as the inhibition of bacteria growth in 50% of the droplets]

For each bacteria density we fit the positive droplet fraction data by the Gompertz function^23,24^. Then we calculate the MIC as the antibiotic concentration value where Gompertz fit crosses the 0.5 viability fraction in droplets (Fig 2G). See further explanation of calculations in Supplementary part 8.

> [We see strong inoculum effect as the MIC rises with the increase of bacteria density (Fig 2H)]

This qualifies as the IE by definition as the increase of MIC is more than 8-fold over the inoculum density increase of two orders of magnitude^13^. This was predicted as similar findings have been reported with the same bacteria strain^20^. We also observed similar IE with different microfluidic setup using smaller droplets and different growth medium^25^. Interestingly, if we look at the growth inhibition *per capita* we see that the amount of cefotaxime needed to suppress the growth remains quite stably around 0.5 μg/CFU. We call this value Minimum Inhibitory Amount (MIA) and suggest including this parameter in future assays describing the antibiotic susceptibility. With limited beneficial interaction between bacteria during antibiotic exposure the MIA remains stable regardless of bacteria inoculum density. However, in case there is a significant increase in MIA, it would hint for a synergistic effect taking place in fighting against antibiotics.

> [Extra advantage of encapsulating bacteria in droplets is that the assay becomes insensitive to fluctuations in inoculum density below encapsulation rate λ ∼0.1]

In these conditions the inoculum density used for preparing the droplets does not have to be controlled precisely as the effective density of bacteria in droplets will always be 1. In our 2nL setup this means that we “lock” the assay into standard 5×10^5^ CFU/mL density. This definitely opens up new ways to design novel “inoculum density resistant” antibiotic susceptibility assays.

> [Unexpectedly we found that droplet-based system is an excellent tool for high-throughput analysis of bacteria clumping]

First, we noticed clumping in control “virtual array” revealed as high fluorescence intensity near value 1000 and above (Fig2E). Clumping is manifested as localization of high intensity pixels in our confocal microscope raw data images (Fig 4A). We measured the relative size of bacteria clumps in droplets by dividing the area of clump with droplet area (more details in Supplementary part 9).

The clumping was extremely intensive in certain fraction of droplets, indicated on Fig 3A as 90th percentile of clump sizes in droplets. In our following analysis we use the 90^th^ percentile value (clump size ∼1% of the droplet) as a threshold for intensive clumping. In most cases we observed a single dominant clump inside the droplet. The true number of clumping events and clump sizes may differ from measured experimental values as the analysis method was not optimized for such analysis (we were not looking for this), and we may have missed some of the clumping events. However, this potential mis-representation is systematic and does not affect the overall findings and their trends. See Supplementary part 3 for explanation.

> [With reanalysis of inoculum density data we saw that clumping of bacteria is modulated by cefotaxime and it is highest at sub-inhibitory drug conditions]

Clumping is most extensive near inhibitory concentrations of cefotaxime both in clumping events and in the average size of clumps (Fig 3BCD). The clumping is also modulated by inoculum density: at low antibiotic concentration the low-bacteria density droplets show more clumping compared to high-bacteria density droplets (bottom section of heat-maps on Fig 3CD). The full results of the clumping events and clump sizes in are shown in Supplementary part 9. Recently, we observed similar clumping trends in smaller droplets using different growth medium^25^.

> [Clumping is considered being the early-phase of biofilm formation]

It seems that clumping (or agglomeration) is a phenotypical growth mode for *E. coli* and cefotaxime increases clumping at sub-MIC concentrations. Similar increased agglomeration at subMIC levels has been described before with beta-lactams and aminoglycosides during the early formation of biofilm in different bacteria^26,27^. We think that our droplet assay has captured the early stages of biofilm formation that is modulated by cefotaxime. In here we use water-in-oil droplets where the stability of the droplets over time is assured by added surfactant. It means that this system does not have any solid surfaces or crevices where bacteria/clumps can attach and form biofilm. This is potentially advantageous in further research into clumping or early-biofilm as the bacteria remain in clump stage.

> [This is the first time such process and its modulation is analyzed in high throughput]

Traditional microscopy and well-plate systems would not have been able to witness this with such high-throughput fashion. As biofilm is one of the key mechanisms in bacterial drug resistance^28^, this will definitely expand investigative horizons in studies regarding biofilm and its formation.

Short conclusion
- Droplet encapsulation enabled us for the first time to quantify the heteroresistance pattern and show the probability distribution of MIC in isogenic bacteria population. Such measurable knowledge of heterogeneity is crucial in understanding how sub-populations of bacteria manage to survive and potentially accumulate resistance increasing mutations.
- We also saw that the relative amount of drug needed to stop bacteria growth is quite stable over wide range of inoculum densities and near these inhibiting conditions the bacteria increasingly clump together. The droplet technology has high potential for high-throughput investigation of early-biofilm formation and its prevention, which is also very important from drug resistance perspective.

## Methods

### Bacteria

In this work we used *Escherichia coli* Dh5α strain that had two plasmids (a very kind gift from Professor Jeff Gore, MIT, US). The first plasmid carries constitutively expressed YFP gene and we used 50 μg/mL of piperacillin for selection. The second plasmid carries TEM-20 gene and we used 50 μg/mL of kanamycin for selection. All experiments, both in bulk and in droplets, were carried out in LB-Lennox media (Roth, Germany). Antibiotic susceptibility experiments were done with cefotaxime. All antibiotics were from Sigma-Aldrich, Germany. Overnight cultures were diluted to required densities with fresh media before the droplet experiments and kept at +4C° until droplet encapsulation.

### Microfludics

The fabrication of the microfluidic chips used in this work has been described before^14,29^. We used separate chips for droplet generation and their fluorescence analysis and more detailed information about their layout can be found in Supplementary part 1. We used Novec HFE-7500 fluorocarbon oil (3M, US) with 2% PFPE-PEG-PFPE surfactant (synthesized according to the protocol by Holtze *et al*.^30^. To control the flow of the oil and reagents in the microfluidic experiments we used rotAXYS positioning system and neMESYS syringe pumps (both from Cetoni, Germany). Droplets were generated with ∼800Hz and analyzed with ∼400Hz frequency. We used conventional 1.5 μL test tubes for off-chip incubation at 37C°. We used labelled dextran-conjugates in our virtual array setup to label different bacteria densities. Cascade Blue™ and Alexa Fluor 647™ were both from Thermo Fisher Scientific, US.

### Fluorescence measurement and data analysis

We measured the fluorescence of droplets in a droplet reading chip, which was mounted on the stage of an A1R confocal microscope (Nikon, Japan). The excitation and detection settings were following: Cascade Blue (405 and 450/50 nm), YFP (488 and 525/50), Alexa 647 (641 and 700/50). Raw images were analyzed with software accompanying the confocal microscope and results were exported as .txt files. Further data analysis was carried out with either MS Office Excel (Microsoft, USA) with Real Statistics Resource Pack (http://www.real-statistics.com/), or with custom made LabVIEW (National Instruments, US) scripts. Droplet signals stand for the peak relative fluorescence intensities allocated to each droplet. See Supplementary parts 3 and 4 for further information about the analysis.

### Virtual array

We made 16 two-fold dilutions of the *E. coli*. First dilution was prepared by refreshing the overnight culture of bacteria in fresh media at 1:10 ratio. The sample loading and droplet generation were described previously in details by Scheler *et al*^14^. In brief, we aspirated 3 uL of each color-coded bacteria dilution into microfluidic tubing spaced with equal volume of oil plugs. Next we compartmentalized the plugs into ∼2 nL droplets (the exact size distribution of droplets is shown in SI-2) and pooled them together as a “virtual array” in a standard 1.5 mL test tube for incubation at 37 °C (Fig 2B) After incubation we analyzed the droplet signals of the virtual array in three different fluorescence detection channels. (Fig 2C)

We position droplets in “virtual array” matrix based on their Cascade Blue and Alexa 647 signal intensities. Then we identify each cluster and determine the bounding box encapsulating the points within and assign color-code number to each gated droplet cluster. The color-coding histogram on Fig 2C shows data from ∼22000 droplets (∼1400 per each color-coded bacteria dilution). In our virtual array experiment we successfully color-coded more than 97% of droplets (See Supplementary part 4 for more details).

### MIC calculation

In experiments with color codding the normalized fraction of positive droplets has dependence qualitatively similar to the behavior shown in Fig. 1C from the main text: the normalized fraction almost always decreases with the antibiotic concentration. For each bacterial density we fit the data by the Gompertz function^23,24^, which is a 2-parameter function:

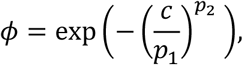

where *c* is the concentration of antibiotic (argument of Gompertz function), *p*_1_ is the value of concentration at which the highest drop of *ϕ* is observed, and *p*_2_ determines the slope of Gompertz function at *c* = *p*_1_. The parameters and their errors are determined by the least square method. We determine MIC by the concentration for which *ϕ* = 1/2, that is, *c_MIC_* = *p*_1_(log 2)^1/*p*_2_^. We estimate error of *c_MIC_* by a minimal value among the error obtained by the error propagation formula applied for *c_MIC_* = *p*_1_ (log 2)^1/*p*_2_^ or the difference between concentrations of the antibiotic closest to *c_MIC_*.

## Acknowledgements

We are very grateful for Professor Jeff Gore from MIT for sharing the bacteria strains.

This research received support from the Foundation for Polish Science within the Team-Tech/2016-2/10 program. P.G. acknowledges support from the Polish National Science Centre based on decision number DEC-2014/12/W/NZ6/00454 (Symfonia). O.S. was supported by the Estonian Research Council grant PUTJD589. O.S. and O-P.S. were supported by the “TTÜ development program 2016– 2022”, project code 2014-2020.4.01.16-0032. W.P. received funding through financial support of the doctoral scholarship from the Polish National Science Centre, scholarship code UMO-2018/28/T/ST4/00318. This project was partially performed in the laboratories funded by NanoFun POIG.02.02.00-00-025/09.

## Author contributions

O.S. conceived the study, designed the research, analyzed data and was responsible for performing the experiments

K.M. was responsible for the data analysis

P.R.D. assisted in data analysis

M.H. assisted in data analysis, was responsible for writing LabView scripts

A.R. assisted in microbiology related work

N.P. assisted in droplet microfluidic experiments

K.S. assisted in confocal microscope measurements

O-P.S. assisted in data analysis

W.P. participated in research design

P.G. conceived the study and assisted in designing the research

O.S., K.M. and P.G. wrote the paper with input from all of the co-authors

## Additional information

Supplementary information is available online

