## Supplementary file for "“Antibiotic inhibition of bacteria growth in droplets reveals heteroresistance pattern at the single cell level”"

1. **Microfluidic chips**

Schematics of i) droplet generation and ii) droplet counting chip. Red scale bar on figures is 1 cm. Schematics on the right are representative and do not show exact size or proportions of the geometries. The exact CAD files are available upon request from the authors.

i. Droplet generation chip

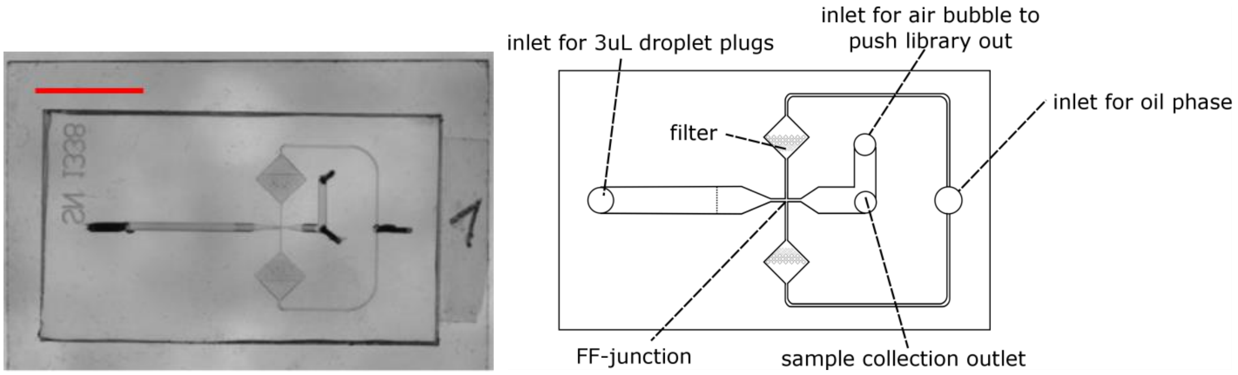

Channels have following dimensions (width x height):

- Main inlet channel: start w:800 x h:800 µm, end at vertical bar position 100x120 µm
- Outlet channel: 800x800 µm
- Oil delivering channel: 200x200 µm
- Flow-focusing junction: 100x120 µm

ii. Droplet counting chip

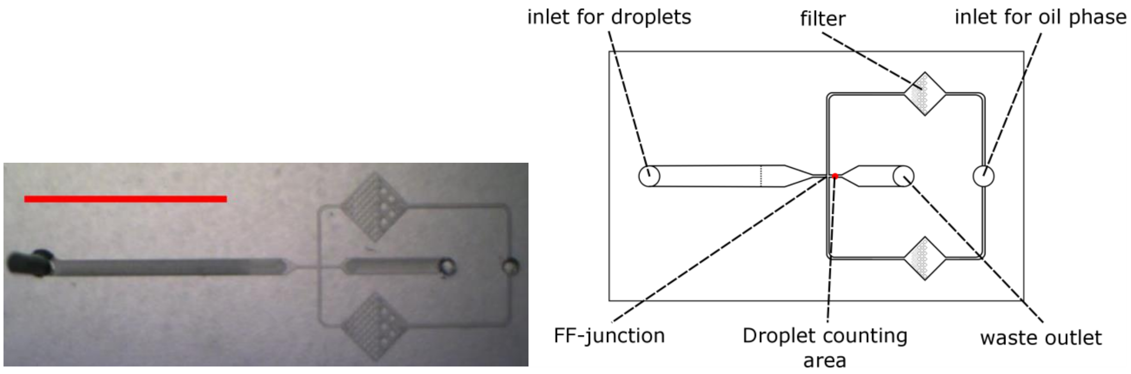

Channels have following dimensions (width x height):

- Droplet inlet channel: 1200x1200 µm at inlet, 170x140 at FF-junction (linear depth change until vertical dashed line)
- Flow-focusing junction: 170x140 µm
- Secondary oil delivering channel: 170x140 µm

### 2. **Droplet size and dispersity**

For measuring the size dispersion of the droplets that were generated from 3μL plugs, we recorded the droplet formation using Photron Fast-Cam 1000K (Japan) camera that was mounted on Nikon SMZ1000 stereoscope (Japan). We measured the whole droplet formation process of a 3μL plug. We picked randomly four different plugs from a dilution series.

| | **Average of 4 tests** | Test1 | Test2 | Test3 | Test4 |
|---|---|---|---|---|---|
| Average droplet volume [nL] | **2.03** | 1.99 | 1.99 | 2.19 | 1.97 |
| Number of droplets generated | **1477.75** | 1505 | 1508 | 1372 | 1526 |
| Coefficient of variation | **5.74%** | 5.09% | 6.06% | 5.35% | 6.44% |

We measured the area of each generated droplet in Image J software and used this data to calculate the volume of each droplet according to the model described previously.[1–3] Briefly, the volume can be calculated by the following formula: $V=(\pi/12)[2D^3-(D-h)^2(2D+h)]$, where D is the diameter of a droplet and h is the height of the channel.

3. **Confocal microscopy data acquisition**

All confocal microscopy experiments were performed using a Nikon A1R setup, based on an inverted confocal microscope Nikon Eclipse Ti-E, equipped with the LU4A laser unit and four-channel detector unit (all by Nikon, Japan). A Nikon Plan Fluor 10x, NA=0.3 objective was used. To maximize the throughput of the system, we acquired fluorescence data using a resonant scanner in a bi-directional single-line mode, with scanning path perpendicular to the microfluidic channel. This allowed for a recording speed of 15360 lines per second (pixel dwell of 0.1 µs, line time of around 65 µs).

At 488 nm, the width of the beam at the focus plane (calculated as doubled 1/e2 radius) was around 1.4 µm. Taking the length of an average droplet in the channel of 145 µm and an average of 40 line scans across each droplet during its passage through the detection zone, we obtain a 37% coverage of the total cross-sectional area of a droplet with the scanning beam (see figure below).

With a widely open pinhole (115 µm), excitation wavelength of 488 nm and a numerical aperture of the objective of 0.3, the thickness of the optical section (i.e. the axial dimension of the detection area) was 57 µm, as reported by the NIS software. This is 41% of the total height of the microfluidic channel. Thus, we can estimate that around 15% of the total volume of the droplet was effectively sampled during the fluorescence data acquisition. Importantly, the scanned lines were distributed evenly along the length of the droplets and the focus plane was set in the center of the channel height. Therefore, we can assume that the data we recorded for each droplet was representative for its whole volume.

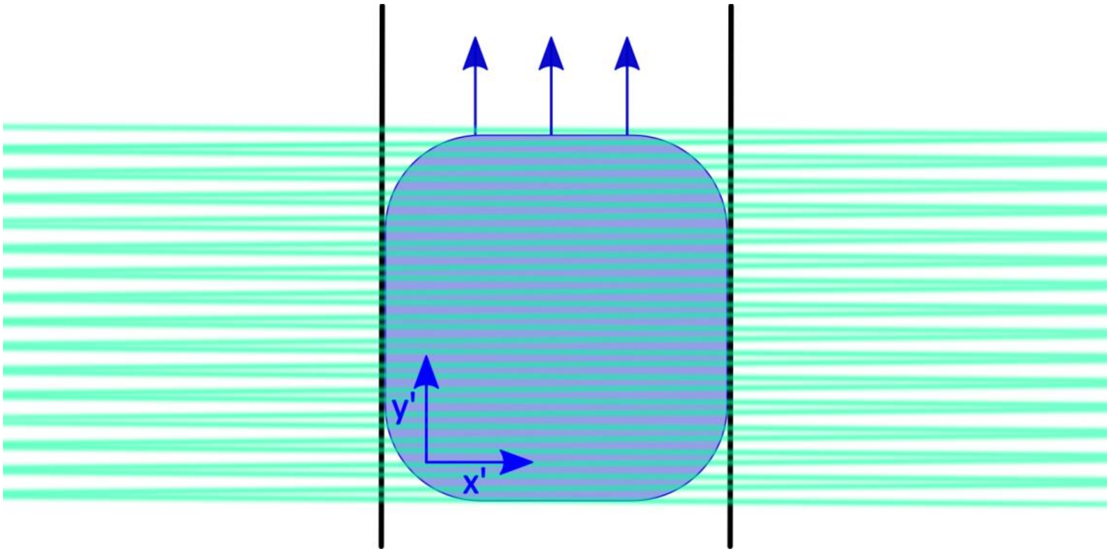

*During data acquisition in line mode, the laser beam is only scanned perpendicularly to the channel (along the x axis). However, the droplet is constantly moving in the y direction. Therefore, in a droplet-bound frame of reference (x',y'), this situation is equivalent to the laser beam scanning through consecutive sections of the droplet in a zig-zagging manner – as schematically depicted in the figure.*

4. **Color-coded droplet addresses in virtual array**

Firstly, raw image stack time-lapse .nd2 files were analyzed with software accompanying the confocal microscope and results were exported as .txt files. Next, the files were analyzed with custom-made LabVIEW script to assign each droplet i) color-code address (Cascade Blue and Alexa Fluor 647 channels) and ii) bacteria fluorescence intensity (YFP channel). The LabVIEW script is available from the authors upon request. In brief, the script plots color-codes in 2D space after which the droplets are assigned addresses according to their clustering into groups based.

Below there are plots showing the virtual arrays according to Cascade Blue (Ch1) and Alexa Fluor 647 (Ch2) intensities. Square boxes mark the gating conditions for each color-code calculated by LabVIEW script. Color-codes from 1 to 16 represent bacteria densities from highest to lowest, respectively. Only droplets that fall within gated boundaries are used for further data analysis. Non color-coded plugs with pure media were added to the beginning and the end of plug train during droplet generation to minimize possible flow rate fluctuations. These droplets locate as color-code 17 in the bottom left and they are omitted from further analysis.  Roman numerals I to XIV stand for different virtual arrays with increasing cefotaxime concentration from 0 to 1024 μg/mL, respectively.

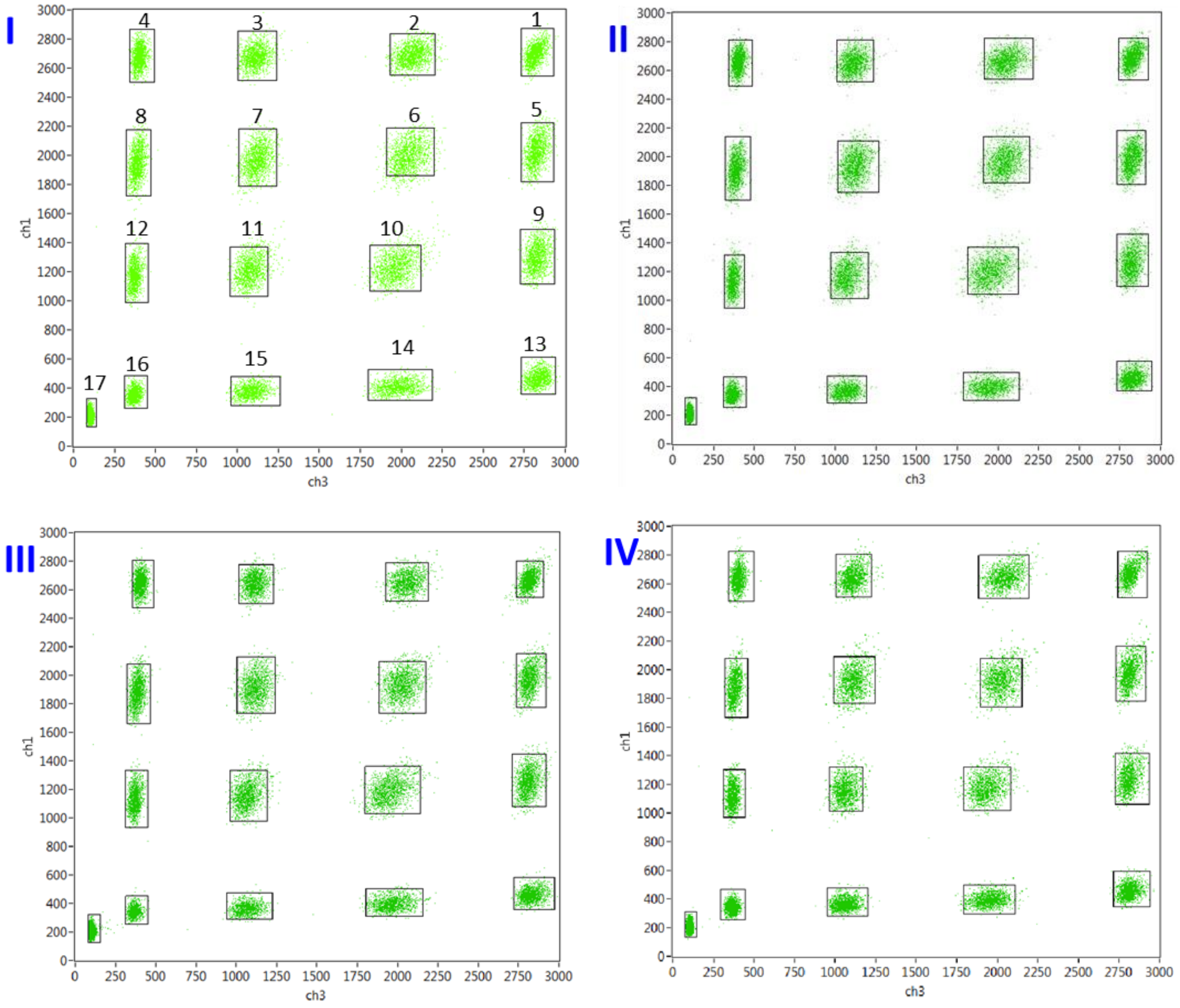

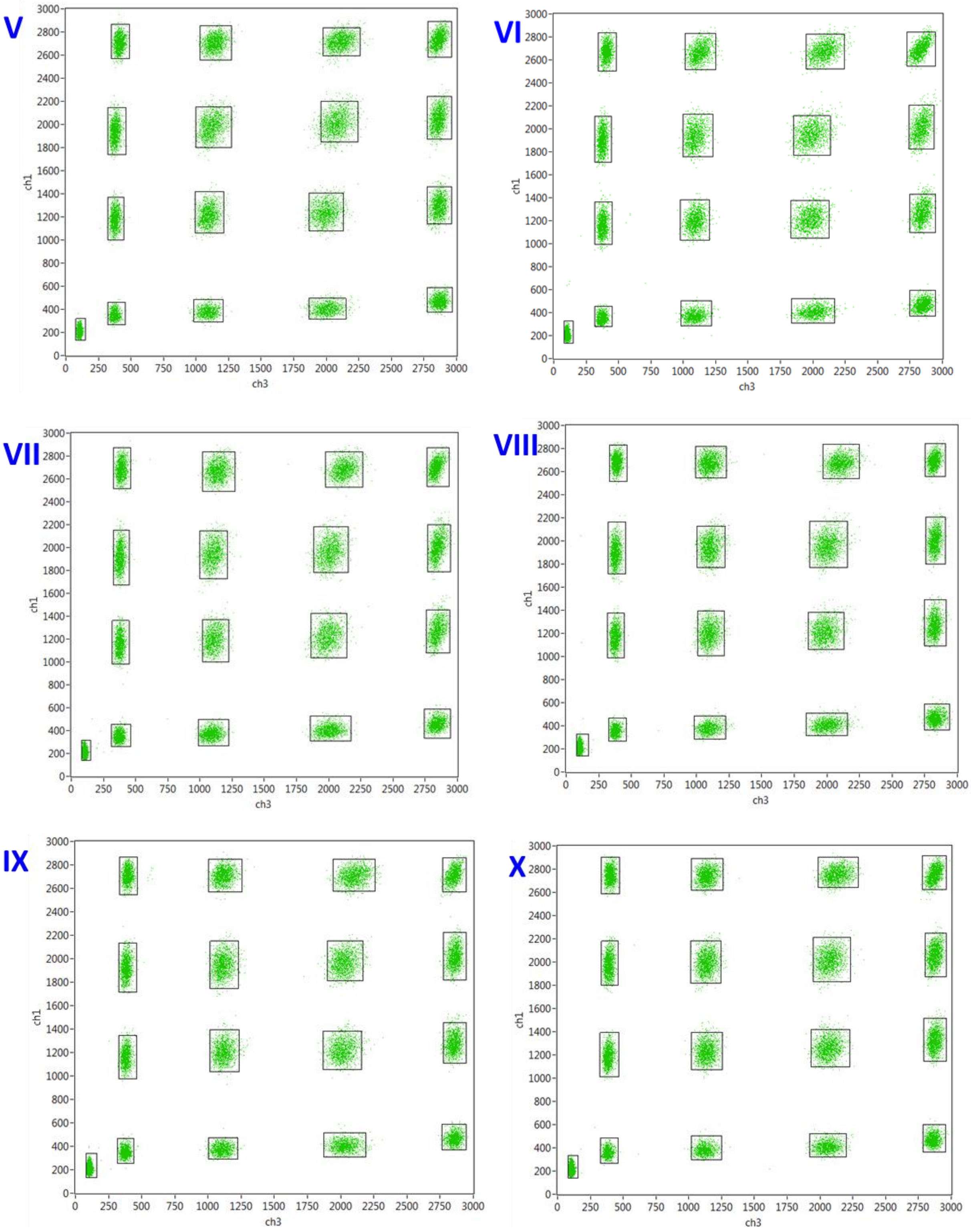
V
VI
VII
VIII
IX
X
ch1
ch3

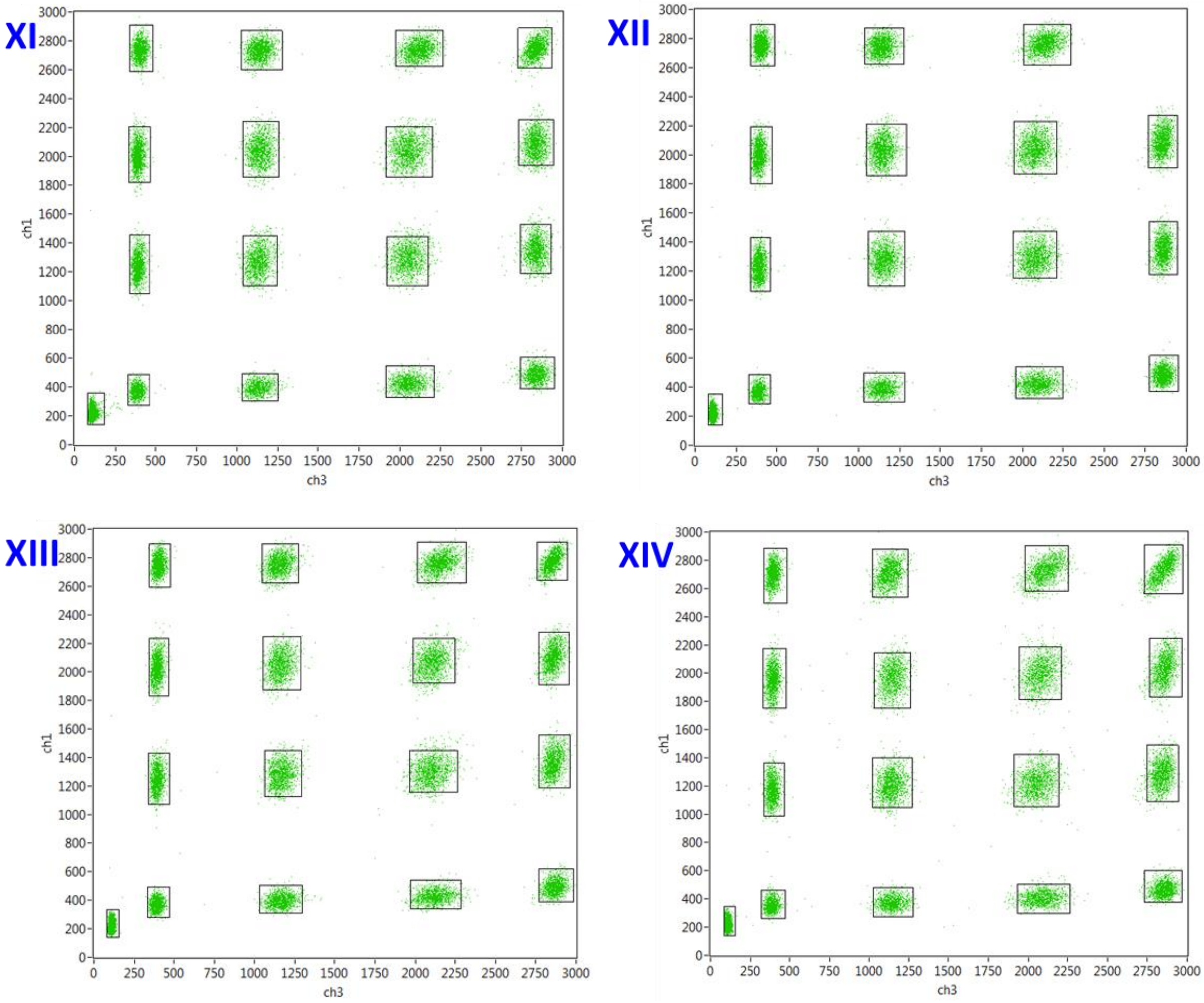

We note that color-code 1 is missing from virtual array XII due to the human error during droplet generation. However, this does not affect the scientific conclusions drawn in the manuscript.

| Virtual array | Cefotaxime [µg/mL] | Droplets in virtual array | AVG droplets in color-code | CV | gated droplet fraction |
|---|---|---|---|---|---|
| I | **0** | 22445 | 1402.8 | 7.35% | 97.51% |
| II | **0.25** | 22094 | 1380.9 | 8.64% | 97.67% |
| III | **0.5** | 22012 | 1375.8 | 10.74% | 97.45% |
| IV | **1** | 22322 | 1395.1 | 5.58% | 97.34% |
| V | **2** | 21623 | 1351.4 | 14.96% | 97.57% |
| VI | **4** | 21970 | 1373.1 | 9.04% | 97.48% |
| VII | **8** | 22952 | 1434.5 | 6.13% | 98.19% |
| VIII | **16** | 23187 | 1449.2 | 11.28% | 98.13% |
| IX | **32** | 21791 | 1361.9 | 8.58% | 98.29% |
| X | **64** | 23376 | 1461.0 | 13.61% | 98.57% |
| XI | **128** | 21763 | 1360.2 | 12.02% | 97.33% |
| XII | **256** | 22018 | 1467.9 | 10.24% | 98.45% |
| XIII | **512** | 22204 | 1387.8 | 4.84% | 97.53% |
| XIV | **1024** | 21493 | 1343.3 | 8.53% | 97.33% |
| | **AVG** | **22232.14** | **1396.1** | **9.40%** | **97.78%** |

Table showing the number of droplets analyzed per each virtual array with average number of droplets/color code shown. Last column shows the percentage of droplets that was gated and assigned with color-code address during the analysis.

5. **Number of bacteria inside droplets**

In our experiments we first emulsify a sample with bacteria and then we incubate droplets obtained by the emulsification. After the incubation, we scan the sample to measure how many droplets we have, $N$, and the number of droplets which give a positive signal, $N_+$. The fraction of positive droplets is denoted by, $f_+ \equiv N_+/N$. In further analysis we assume that in each droplet there is a random number of bacteria after the emulsification. Therefore there is the probability $f_+$ that a droplet gives a positive signal (i.e. contains at least one bacteria after the emulsification) and the probability $1 - f_+$ that a droplet gives a negative signal (contains no bacteria after the emulsification). From this perspective our experiment with droplets relies on performing of $N$ Bernoulli trials with the probability of success equal to $f_+$. It follows from the central limit theorem that in the limit of large number of droplets (i.e. large number of Bernoulli trials) the fraction of positive droplets have a Gaussian distribution with the average $f_+$ and the dispersion $\sigma\_f\_+ = \surd(f_+ (1 - f_+)/N)$. The central limit theorem cannot be applied for the case when $f_+ = 0$ or $f_+ = 1$. In this case we estimate error of the measurement of $f_+$ using Bayesian inference by, $\sigma_{f_+} = 1/N$. The fraction of positive droplets can be used to calculate the average number of bacteria inside a droplet, $\langle N_{CFU} \rangle \equiv N_{bact}/N$, where $N_{bact}$ denotes number of bacteria inside all droplets. It is possible after assumption that bacteria are closed in droplets according to Poisson statistics.[4–6] It follows that $\langle N_{CFU} \rangle (f_+) = -\log(1 - f_+)$. For the interpretation of our data we use the average number of bacteria inside droplets which gives a positive signal, $\langle N_{CFU} \rangle_+ \equiv N_{bact}/(N - N_+)$. In terms of the fraction $f_+$ it is given by the following formula, $\langle N_{CFU} \rangle_+(f_+) = -[\log(1 - f_+)]/f_+$. The above formulas are used to determine $\langle N_{CFU} \rangle (f_+)$ and $\langle N_{CFU} \rangle_+(f_+)$ in our experiments shown in Fig. 1 from the main text. Their errors are determined from the error propagation formula [Introduction to error analysis, the study of uncertainties in physical measurements, John Taylor] and the error of $f_+$, i.e. $\sigma_{f_+}$, described in this paragraph. The above two expressions for $\langle N_{CFU} \rangle (f_+)$ and $\langle N_{CFU} \rangle_+(f_+)$ lead to the relation $\langle N_{CFU} \rangle_+ = \langle N_{CFU} \rangle / \left(1 - exp(-\langle N_{CFU} \rangle)\right)$.

6. **Initial bacteria density in different dilutions**

The above formula for average number of bacteria in droplets cannot be used when all droplets give a positive signal. It happens in some of our 16 dilutions. To calculate density in all dilutions, we assessed first only the libraries that contained less than 100% positive droplets (libraries 6-16). For these libraries we determined $\langle N_{CFU} \rangle$ values with errors using the method described in the previous paragraph. As the consecutive libraries were serially diluted, the estimated numbers of bacteria provide also the dilution ratio between the libraries, $\langle N_{CFU} \rangle_i = n_1/x^{i-1}$. We calculate $n_1$ and $x$ parameters by fitting this curve to the numbers of bacteria (and their errors) obtained for the libraries 6-16 using least squares method. The estimated dilution ratio between the libraries was equal to $x = 2.23 \pm 0.03$ and $n_1 = 340 \pm 40$ which we use to calculate densities in all samples. For each experiment we also determine $\langle N_{CFU} \rangle_+$ from $\langle N_{CFU} \rangle$ (and its error from the error propagation formula), as described at the end of the previous paragraph. The result of the calculations are presented in accompanying figure ad table.

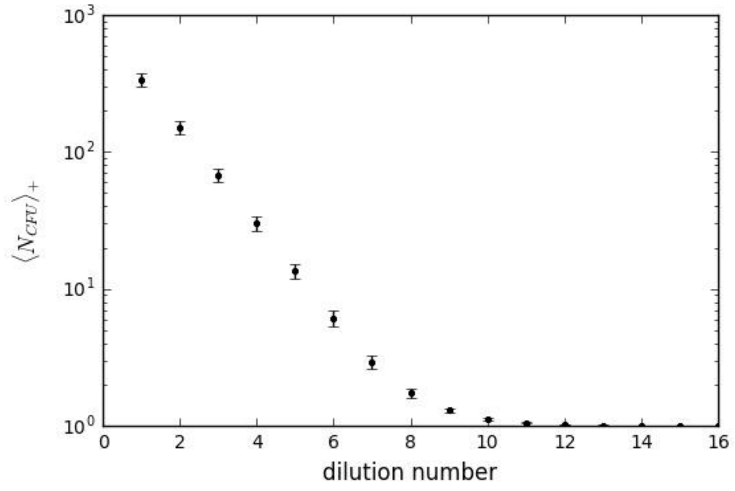

| Library | NCFU+ | error |
|---|---|---|
| 1 | 337,54 | 37,43 |
| 2 | 151,19 | 16,92 |
| 3 | 67,72 | 7,77 |
| 4 | 30,34 | 3,62 |
| 5 | 13,59 | 1,71 |
| 6 | 6,10 | 0,80 |
| 7 | 2,92 | 0,34 |
| 8 | 1,73 | 0,13 |
| 9 | 1,30 | 0,05 |
| 10 | 1,13 | 0,02 |
| 11 | 1,06 | 0,01 |
| 12 | 1,02 | 0,00 |
| 13 | 1,01 | 0,00 |
| 14 | 1,00 | 0,00 |
| 15 | 1,00 | 0,00 |
| 16 | 1,00 | 0,00 |

### 7. Individual MIC distribution in the population

As described in the main text, the probability distribution if individual MIC is related to the normalized fraction of positive droplets, $\int_c^\infty p(m)dm = f_+(c)/f_+(0)$. Taking derivative of the this formula gives, $p(c) = -\frac{d}{dc} f_+(c)/f_+(0)$. To calculate the derivative we use the data points from Fig. 1C for different concentrations $c_i$ according to the following formula $p((c_{i+1} + c_i)/2)$=$\left(\frac{f_+(c_{i+1})}{f_+(0)} - \frac{f_+(c_i)}{f_+(0)}\right)/(c_i - c_{i+1})$. Errors of the above probability distribution are determined by error propagation formula from errors of $f_+(c_i)$, $f_+(c_{i+1})$ and $f_+(0)$.

### 8. MIC determination and inoculum effect

In experiments with color codding the normalized fraction of positive droplets has dependence qualitatively similar to the behavior shown in Fig. 1C from the main text: the normalized fraction almost always decreases with the antibiotic concentration. For each bacterial density we fit the data by the Gompertz function [7,8], which is a 2-parameter function:

$$\phi = \exp\left(-\left(\frac{c}{p_1}\right)^{p_2}\right),$$

where $c$ is the concentration of antibiotic (argument of Gompertz function), $p_1$ is the value of concentration at which the highest drop of $\phi$ is observed, and $p_2$ determines the slope of Gompertz function at $c = p_1$. The parameters and their errors are determined by the least square method. We determine MIC by the concentration for which $\phi = 1/2$, that is, $c_{MIC} = p_1(\log 2)^{1/p_2}$. We estimate error of $c_{MIC}$ by a minimal value among the error obtained by the error propagation formula applied for $c_{MIC} = p_1(\log 2)^{1/p_2}$ or the difference between concentrations of the antibiotic closest to $c_{MIC}$.

9. **Bacteria clumping analysis**

In our experiments we observed clumping of bacteria manifested as clusters of high-intensity pixels in confocal line-scan stack (**red arrow** in B). First, we noticed clumping in droplets of control “virtual array” without the addition of antibiotics. To quantify the clumping events in our experiments we measured i) the existence of such high-intensity pixels in each droplet and ii) the relative size of each cluster if such event occurred in droplet.

i) Clumping events were analyzed using object count function in confocal software with following settings (Fig C). Thresholds: low (3000, threshold for high-intensity pixel) and high (4095, maximum signal), Smooth: OFF, Clean: OFF, Fill holes: ON, Separate: OFF. This measurement gives also shows the area of each clump ($A_c$)

Relative size of clumps (S) was measured as area of clump ($A_c$) / area of droplet ($A_d$). Droplet sizes ($A_d$) were measured using binary function with following settings (Fig D). Thresholds: low (100, minimum signal) and high (4095, maximum signal), Smooth: 4x, Clean: 2x, Fill holes: ON, Separate: OFF.

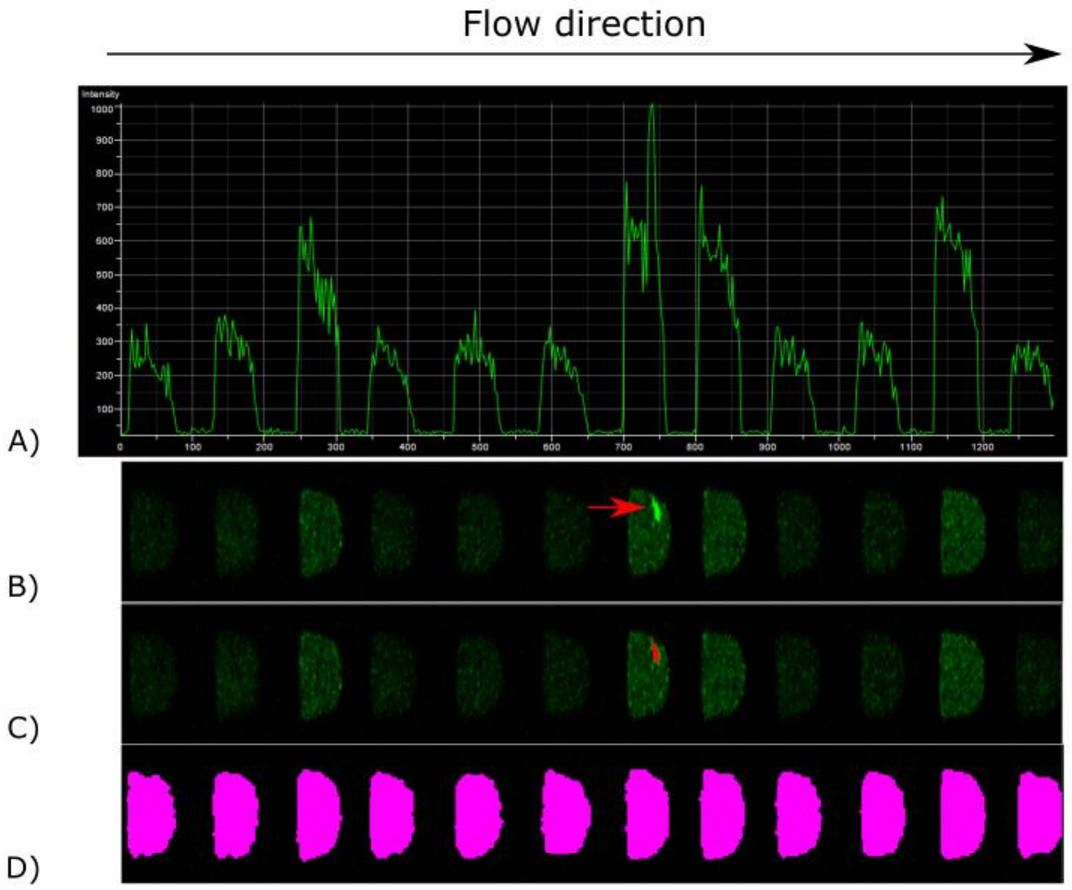

i) We decided to measure clump size as relative to droplet size because the flow speed of droplets during imaging can fluctuate as the population of droplets pushed through the chip decreases. Such flow fluctuation can cause small changes in imaged droplet areas.

ii) Clumping is not caused by debris or dust in the growth media as there were no clumping events detected in the beginning of experiment at 0h before the incubation.

iii) In case of clumping the droplet is usually dominated by a single dominant clump. We observed that with ~80% of the clumping events the size of the biggest clump ($A_c$) was more than half of the total area of measured clumps ($A_{sum}$). The dominance of single clump was similarly near 80% in all bacteria inoculum densities (**red dots** on the graph). **Blue dot** shows the average of red dots with respective standard deviation as error bars.

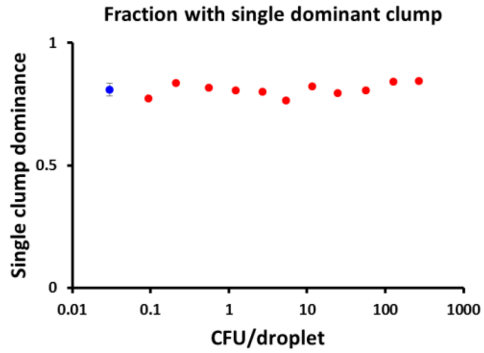

iv) The clumping was extremely intensive in certain fraction of droplets and this fraction is shown on the figure as top 90 decile of clump sizes in droplets. In our following analysis we use the 90th decile value (clump size ~1.06%) as a threshold for “intensive clumping”.

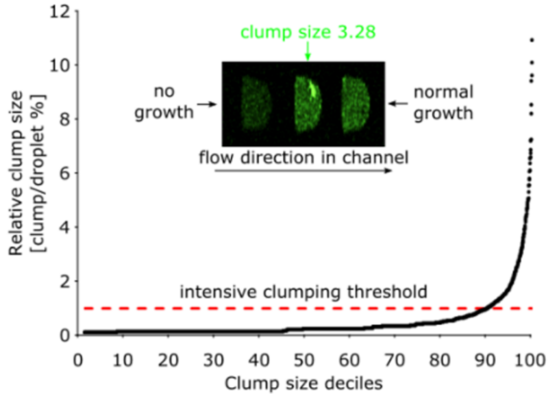

v) The same analysis criteria were then used to measure clumping in virtual array experiments with different antibiotic concentrations.

vi) Following table shows the data about the percentage of droplets where bacteria demonstrated intensive clumping.

**Percentage of droplets with extensive clumping**

| | | | | | | | | | | | | |
|---|---|---|---|---|---|---|---|---|---|---|---|---|
| **Cefotaxime [ug/mL]** | | | | | | | | | | | | |
| **1024** | | | | | | | | | | | 6,67% | |
| **512** | | | | | | | | | | | 22,22% | |
| **256** | | | | | | | | | | | | |
| **128** | | | | | | | | | | | 28,97% | |
| **64** | | | | | | | | | | 7,77% | 42,66% | |
| **32** | | | | | | | | | 15,79% | 18,51% | 7,40% | |
| **16** | | | | | | | | 0,00% | 18,07% | 29,36% | 0,82% | |
| **8** | | | | | | | 0,00% | 10,31% | 26,93% | 2,52% | 0,25% | |
| **4** | | | | | 6,67% | 2,62% | 5,88% | 11,94% | 2,75% | 0,75% | 0,27% | |
| **2** | | | | 6,38% | 3,91% | 4,92% | 8,99% | 8,78% | 2,36% | 1,27% | 0,46% | |
| **1** | 8,70% | 2,94% | 5,49% | 5,28% | 4,82% | 5,50% | 8,36% | 7,66% | 3,13% | 1,59% | 0,70% | |
| **0,5** | 0,77% | 2,81% | 3,91% | 3,23% | 4,58% | 4,95% | 6,73% | 3,70% | 1,38% | 0,36% | 0,14% | |
| **0,25** | 5,23% | 3,69% | 3,98% | 3,74% | 3,45% | 5,15% | 7,30% | 5,41% | 2,57% | 0,93% | 0,19% | |
| **0** | 2,50% | 3,15% | 2,82% | 4,75% | 3,45% | 4,31% | 4,26% | 3,73% | 1,47% | 1,07% | 0,19% | |
| | **1,06** | **1,13** | **1,30** | **1,73** | **2,92** | **6,10** | **13,59** | **30,34** | **67,72** | **151,19** | **337,54** | **bacteria density [CFU/positive droplet]** |

vii) Following table shows the average relative size of clumps in droplets, calculated as the percentage of droplet area. Relative size of clumps (S) was measured as area of clump ($A_c$) / area of droplet ($A_d$).

**Average clump size in droplets**

| | | | | | | | | | | | | |
|---|---|---|---|---|---|---|---|---|---|---|---|---|
| **Cefotaxime [ug/mL]** | | | | | | | | | | | | |
| **1024** | | | | | | | | | | | 1,80% | |
| **512** | | | | | | | | | | | 2,98% | |
| **256** | | | | | | | | | | | | |
| **128** | | | | | | | | | | | 7,40% | |
| **64** | | | | | | | | | | 8,32% | 3,44% | |
| **32** | | | | | | | | | 4,23% | 6,10% | 1,45% | |
| **16** | | | | | | | | 0,00% | 6,32% | 2,86% | 1,29% | |
| **8** | | | | | | | 0,00% | 6,13% | 5,99% | 1,38% | 1,19% | |
| **4** | | | | | 4,60% | 4,06% | 4,62% | 3,10% | 1,37% | 1,77% | 1,22% | |
| **2** | | | | 3,34% | 3,20% | 5,40% | 4,00% | 2,34% | 1,59% | 1,69% | 1,45% | |
| **1** | 3,51% | 2,59% | 3,94% | 5,70% | 5,26% | 4,83% | 2,92% | 2,15% | 1,59% | 1,47% | 1,41% | |
| **0,5** | 1,63% | 3,47% | 4,38% | 4,52% | 3,47% | 3,06% | 2,62% | 2,03% | 1,50% | 1,59% | 1,17% | |
| **0,25** | 6,29% | 5,20% | 4,16% | 4,31% | 4,01% | 3,11% | 2,41% | 2,02% | 1,60% | 1,43% | 1,26% | |
| **0** | 2,00% | 4,13% | 3,83% | 3,20% | 2,58% | 2,70% | 2,50% | 1,83% | 1,47% | 1,34% | 1,43% | |
| | **1,06** | **1,13** | **1,30** | **1,73** | **2,92** | **6,10** | **13,59** | **30,34** | **67,72** | **151,19** | **337,54** | **bacteria density [CFU/positive droplet]** |

### 10. List of References
